# Integrative Thermodynamics Strategies in Microbial Metabolism

**DOI:** 10.1101/2025.09.09.672066

**Authors:** Martijn Bekker, Oliver Ebenhöh

## Abstract

Microbial metabolism is intricately governed by thermodynamic constraints that dictate energetic efficiency, growth dynamics, and metabolic pathway selection. Previous research has primarily examined these principles under carbon-limited conditions, demonstrating how microbes optimize their proteomic resources to balance metabolic efficiency and growth rates. This study extends this thermodynamic framework to explore microbial metabolism under various non-carbon nutrient limitations (e.g., nitrogen, phosphorus, sulfur). By integrating literature data from a range of species it is shown that growth under anabolic nutrient limitations consistently results in more negative Gibbs free energy (Δ*G*) values for the Net Catabolic Reaction (NCR), when normalized per unit of biomass formed, compared to carbon-limited scenarios. The findings suggest three, potentially complementary hypotheses: (1) Proteome Allocation Hypothesis: microbes favor faster enzymes to reduce proteome fraction used for catabolism, thus freeing proteome resources for additional nutrient transporters; (2) Coupled Transport Contribution Hypothesis: The more negative Δ*G* of the NCR may in part stem from the increased reliance on ATP-coupled or energetically driven transport mechanisms for nutrient uptake under limitation; (3) Bioenergetic Efficiency Hypothesis: microbes prefer pathways with more negative Δ*G* to enhance cellular energy status, such as membrane potentials or ATP/ADP ratio, to support nutrient uptake under anabolic limitations. This integrative thermodynamic analysis broadens the understanding of microbial adaptation strategies and provides valuable insights for biotechnological applications in metabolic engineering and fermentation process optimization.

## 1. Introduction

Microbial metabolism is governed by the fundamental principles of thermodynamics, constraining energy efficiency, growth rates, and metabolic trade-offs [1–7]. Understanding the Gibbs free energy (Δ*G*) requirements of different metabolic pathways is critical for predicting microbial behavior under various environmental constraints.

The framework of mosaic approach in non-equilibrium thermodynamics, as described by Westerhoff et al. [1,2,8], provides a theoretical basis for understanding microbial growth and thermodynamics forces involved. This approach views microbial metabolism in terms of energy fluxes, distinguishing between output flow (catabolic processes) and input flow (substrate utilization). A key insight from this perspective is that microbial systems are often optimized towards maximum growth rate rather than maximum growth yield [1,6,9,10] challenging previous assumptions in microbial bioenergetics. Interestingly, under nutrient limitations, deviations from theoretical ATP yields (*Y*_ATP_) arise due to maintenance processes and/or futile cycles [11–13]. According to Westerhoff [8] these deviations emphasized the need to account for non-growth related maintenance requirements.

Flamholz et al. [9] approached thermodynamic modeling of microbial metabolism by use of the Gibbs free energy equation in combination with the Haldane rate law, which defines the limitations (with respect to forward and reverse *k*_cat_ and *K*_M_) of enzyme characteristics. Using this aspect Flamholz et al. applied the rules for a single enzymatic reaction (i.e. the Haldane equation) to complete metabolic pathways [9]. This allowed explanation of the presence of the Entner-Doudoroff (ED) pathway and its preferential use in some cases over the Embden-Meyerhoff-Parnass (EMP) pathway, which appears counter-intuitive considering that the ED pathway only produces one ATP per glucose converted to pyruvate, in comparison to the two ATP of the EMP pathway. It was eloquently shown that, to obtain the same ATP production rate, the proteome resource requirements for the ED pathway would be 5-fold lower (using measured kinetics) than for the EMP pathway. By producing only one instead of two ATP, the net Δ*G* of the ED pathway is more negative than that of the EMP pathway. This in turn allows for higher *k*_cat_-values of the forward reactions allowed within the limitations of the more negative Δ*G* of the ED pathway as compared to the EMP pathway [9,14], leading to faster ATP production rates with a smaller enzyme investment, despite the twofold lower yield of ATP per glucose converted to pyruvate.

This hypothesis was subsequently expanded to explain overflow metabolism in *Escherichia coli* [6,14–17], the Crabtree effect [18–20] and mixed acid versus homolactic fermentation [21]. By using these concepts it can be hypothesized that all these observations are a consequence of proteome resource allocation constraints [3] imposed by Δ*G* and ATP yield trade-offs [22]. These are of high relevance for a better understanding of the intricacies with which thermodynamics influence the selected metabolic pathways for ATP generation and the consequences of this for proteome resource allocation.

The above endeavors all focus on explaining the change from carbon-limited growth conditions to batch conditions (i.e. carbon-excess conditions) [6,21,23]. None of these studies have focused on non-carbon limited growth conditions. Interestingly, such conditions show a much higher maintenance requirement than carbon limited growth [13,24–26] and calculations on the qATP_m_ indicated higher maintenance especially at low growth rates [24]. However, these observations are not yet included in recent studies on thermodynamics of microbial growth.

The motivation behind this research paper is therefore to extend previous studies with an integrative thermodynamic analysis of published literature of non-carbon limitations and the effects on the thermodynamics of growth. By utilizing this extended theoretical model we aim to widen our understanding of the fundamental thermodynamic phenomena observed under specific nutrient limited conditions [24,25,27].

## 2. Theory

Microbial growth is often conceptually viewed as a thermodynamic energy converter, in which the energetically favorable reactions of catabolism drive anabolism [1,28,29]. The ‘coupling agent’ is the energy currency metabolite ATP, which is produced in catabolic pathways, exploiting the free energy gradient of the conversion of nutrients into catabolic products. This ATP, in turn, drives thermodynamically unfavorable reactions of anabolism (see Figure 1 for a schematic representation). Whereas ATP is a key intermediate and quantities such as *Y*_ATP_, the ATP yield per mole substrate consumed, are important characteristics for the thermodynamic efficiency of growth, these are notoriously difficult to measure and are often inferred using metabolic models and rely on certain assumptions, such as the P/O ratio. We here propose an approach that does not depend on the knowledge of ATP yields or demands, but that instead relies exclusively on knowledge of the overall macrochemical growth equation.

**Figure 1.**
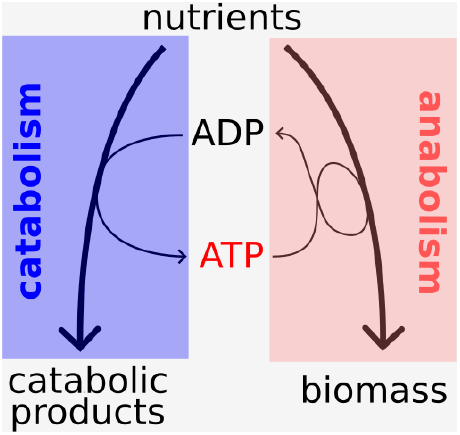
Microbial growth as energy converter

### Box 1: Net Catabolic Reaction

The energy converter model describes microbial growth as the coupling of two processes. The free energy released in catabolism drives the thermodynamically less favorable anabolism. Assume a macrochemical growth equation of

1.78 CH_2_O + 0.15 NH_3_ + 0.73 O_2_ → CH_1.79_O_0.57_N_0.15_ + 0.78 CO_2_ + 1.11 H_2_O,where CH_2_O stands for one carbon mole glucose. Here, *γ*_*S*_ = 4 and *γ*_*X*_ = 4.2, resulting in a theoretical yield of *γ*_*S*_/*γ*_*X*_ = 4/4.2 ≈ 95.2%. This results in the ideal anabolic reaction

1.05 CH_2_O + 0.15 NH_3_ ***→*** CH_1.79_O_0.57_N_0.15_ + 0.05 CO_2_ + 0.38 H_2_O, leaving the **Net Catabolic Reaction**

0.73 CH_2_O + 0.73 O_2_ → 0.73 CO_2_ + 0.73 H_2_O

with a reaction energy of Δ*G*_X/S_ ≈ −350 kJ/C−mol biomass.

An important quantity in the center of our focus is the amount of Gibbs free energy that needs to be released by catabolic processes to produce one carbon mole of biomass. By measuring the catabolic energy requirement to produce one carbon mole of biomass, this quantity provides an important thermodynamic characteristic of microbial growth. We will in the following refer to it as the catabolic Gibbs free energy and denote it with Δ*G*_X/S_.

The quantity Δ*G*_X/S_ is determined from the macrochemical growth equation as follows: First, the macrochemical growth equation is normalized to the production of one carbon mole biomass. Second, the net anabolic reaction to form one carbon mole biomass is subtracted, leaving only the catabolic conversion involved in the formation of one carbon mole biomass. We term the remaining chemical conversion the Net Catabolic Reaction (NCR). In order to define the net anabolic reaction, we consider an idealized process that converts nutrient carbons into biomass with a maximally possible yield. If biomass is less reduced than the substrate, the theoretical carbon yield of this anabolic process is 1 (100%), assuming the presence of external electron acceptors, such as oxygen. If the biomass is more reduced than the substrate, some nutrient carbons need to be oxidized to maintain the redox balance, with a maximal theoretical carbon yield of 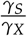, where *γ*_*S*_ and *γ*_*X*_ represent the degrees of reduction of the substrate and biomass, respectively. The NCR can easily be determined from a macrochemical growth equation (see Box 1) and we calculate the associated reaction energy Δ*G*_X/S_ using the eQuilibrator API [9].

It should be noted that the ‘ideal’ anabolic reaction is an abstract concept and there is no guarantee that such a reaction can actually be realized. Interestingly, though, we recently showed with genome-scale metabolic models that feasible flux distributions indeed do exist, which closely approximate (>95%) the theoretical carbon yield, which are fully operable provided sufficient ATP is provided by other pathways [30]. However, whether such idealized pathways actually exist is secondary to our investigation. Instead, a simple, unambiguous definition is important that gives rise to a single calculation method that can be applied to all micro-organisms, carbon sources and conditions. The NCR and the associated catabolic free energy Δ*G*_X/S_ fulfill all these conditions, thus they provide a thermodynamic measure of microbial growth which is comparable across organisms and growth conditions.

## 3. Methods

### 3.1 Net Catabolic Reaction

Every dataset is individual. We therefore first convert every single chemostat dataset so that all measured exchange fluxes are given in the unit C−mol (C−mol biomass)^*−*1^ h^*−*1^ or mol (C−mol biomass)^*−*1^ h^*−*1^ for non carbon containing compounds (in particular O_2_). These values are assembled in a vector, such that consumed compounds are counted negative. For convenience, we further assign biomass to the index 0. For *n* additionally measured exchange fluxes, this results in an *n* + 1-dimensional vector *e* = (*e*_0_, …, *e*_*n*_)^*T*^, which contains the measured exchange fluxes in carbon mole units. For consumed metabolites the corresponding coefficients are negative. If all metabolites were measured and 100% elemental balance were given, these quantities would directly be stoichiometric coefficients of the macrochemical growth equation. Because of imperfect measurements, such a directly determined macrochemical growth equation would not be mass balanced. We therefore fit a mass-balanced macrochemical growth equation to the data in the following way:

1. From the experimentally measured catabolic products we derive a list of net catabolic reactions, which result in the production of the observed metabolite. For example, if acetate was measured for growth on glucose, the assumed net catabolic reaction would be

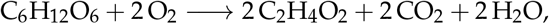

which is further normalized to carbon mole as

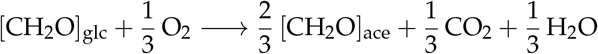

and converted into a vector *v* containing the stoichiometric coefficients. The coefficients of this vector are ordered, such that the first *n* + 1 entries correspond to the experimentally measured metabolites, including biomass. If *k* catabolic products were determined, this results in *k* vectors *v*_1_ … *v*_*k*_.
2. An additional vector *v*_0_ is defined characterizing biomass growth as follows: From the elemental composition of the biomass, X, the degree of reduction *γ*_*X*_ is determined after [31]. The degree of reduction of the carbon source, S, is denoted by *γ*_*S*_. The idealized anabolic equation is then assumed to be

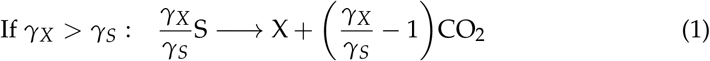

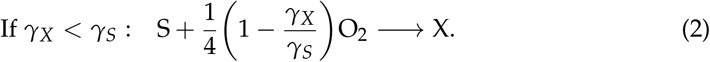

Because compounds containing nitrogen, phosphorus or sulfur have not been systematically measured, these equations only contain the carbon balance.
3. A linear combination is determined that fits the experimental vector *e* best by minimizing the residual squares,

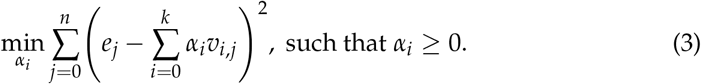

These residuals are reported, e.g., in Table 1. The fitting procedure is performed with in python with the method `scipy.optimize.minimize.`
4. The resulting vector *m*, which, when normalized to one unit of biomass formed, has the coefficients

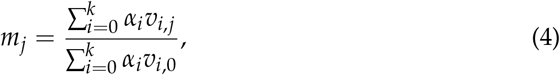

represents the macrochemical growth equation, normalized to the production of one carbon mole biomass, which fits best to the experimentally measured exchange fluxes. The combination of catabolic routes (represented by the vector 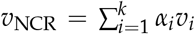 forms the *Net Catabolic Reaction* (NCR), which is used for further calculations.

**Table 1.** Summary of rates and thermodynamic properties under different continuous growth conditions and nutrient limitations of *Saccharmoyces cerevisiae* (data extracted from [25]). The table provides a schematic overview of respiration and fermentation rates (both in C-mol glucose/C-mol biomass/h consumed in the respective pathway) and the calculated overall thermodynamics of the Net Catabolic Reaction (NCR) across various nutrient-limited continuous growth conditions. The quality of fit reports the sum of residual squares of the fitted versus experimental fluxes (measured in C−mol L^*−*1^ h^*−*1^). Higher numbers indicate a carbon recovery coefficient diverging from 100% and, as a result, a poorer fit to the data.

| Condition | Limitation | Respiration | Fermentation | $\Delta G$(kJ/C-mol biomass) | Quality of fit ($r^2$) |
| --- | --- | --- | --- | --- | --- |
| Aerobic | Carbon | 0.7 | 0 | -344 | 0.02 |
|  | Nitrogen | 0.79 | 6.17 | -610 | 3.03 |
|  | Phosphorus | 1.15 | 6.17 | -784 | 5.22 |
|  | Sulfur | 0.77 | 3.41 | -497 | 1.23 |
| Anaerobic | Carbon | 0 | 7.27 | -260 | 2.22 |
|  | Nitrogen | 0 | 10.50 | -376 | 8.73 |
|  | Phosphorus | 0 | 10.50 | -391 | 10.70 |
|  | Sulfur | 0 | 9.65 | -346 | 15.59 |

### 3.2 Net catabolic Gibbs free energy ΔG_X/S_

The idealized anabolic equation (1) or (2) is subtracted from the vector *m*, yielding the vector *v*_NCR_ describing the *Net Catabolic Reaction* (NCR). The vector *v*_NCR_ represents a fully mass balanced chemical conversion that is normalized to the formation of one carbon mole biomass. The corresponding energy of reaction is exactly the net catabolic Gibbs free energy Δ*G*_X/S_ required to produce one carbon mole of biomass. This value is determined using the eQuilibrator API [9], taking environmental data, such as pH and temperature, into account as reported in the original literature. Because in most experiments, extracellular concentrations of nutrients and catabolic products have not been measured, we determine the reaction energies for biochemical standard conditions with concentrations at 1 mmol L^*−*1^. While the actual concentrations will certainly affect the reaction energies quantitatively, this effect can be expected to be rather small. As argued in [30], every order of magnitude deviation from the biochemical standard concentration will result in a shift of *∼*5.7 kJ mol^*−*1^. With catabolic reaction energies of 300 kJ mol^*−*1^ and, in some cases, considerably more, even an uncertainty in concentrations of several magnitudes will result only in a rather small relative error. Whereas it would be interesting to systematically measure concentrations and thus quantify the concentration-dependent effect, including measured concentrations would not qualitatively alter our results and therefore also not affect the conclusions we draw here.

### 3.3. Estimation of errors

The group contribution method underlying the estimation of reaction energies [9,32,33] naturally involves significant uncertainties. However, these uncertainties are systematic and effect all datasets similarly. Because we are interested in comparing catabolic free energies across experiments, organisms and conditions, we decided to ignore this systematic error but rather estimate the error that stems from the measurement uncertainties, usually indicated by a carbon recovery coefficient deviating from 100%. We estimate the error as follows: We compare the fitted substrate uptake rate (in vector *m*) with the experimentally measured rate (in vector *e*) and make two extreme assumptions to ‘correct’ this difference. For simplicity, we assume the fitted uptake rate is smaller than the experimental rate (the argument below works exactly the same, but opposite, if the rate is larger). First, we assume that the missing carbon was completely converted into biomass and add this conversion to the vector *m* and repeat the subsequent calculation. This results in a lower value of Δ*G*_X/S_, because the catabolic free energy is now normalized to a higher biomass production rate. Second, we assume that the missing carbon was completely respired to CO_2_, and add the corresponding respiration reaction to the vector *m*, before performing the subsequent calculations again. This results in a higher value of Δ*G*_X/S_, because the additionally respired carbon will add to the catabolic reaction energy. These errors are given as error bars in Figure 2 and, e.g., reported in Table 2.

**Table 2.**
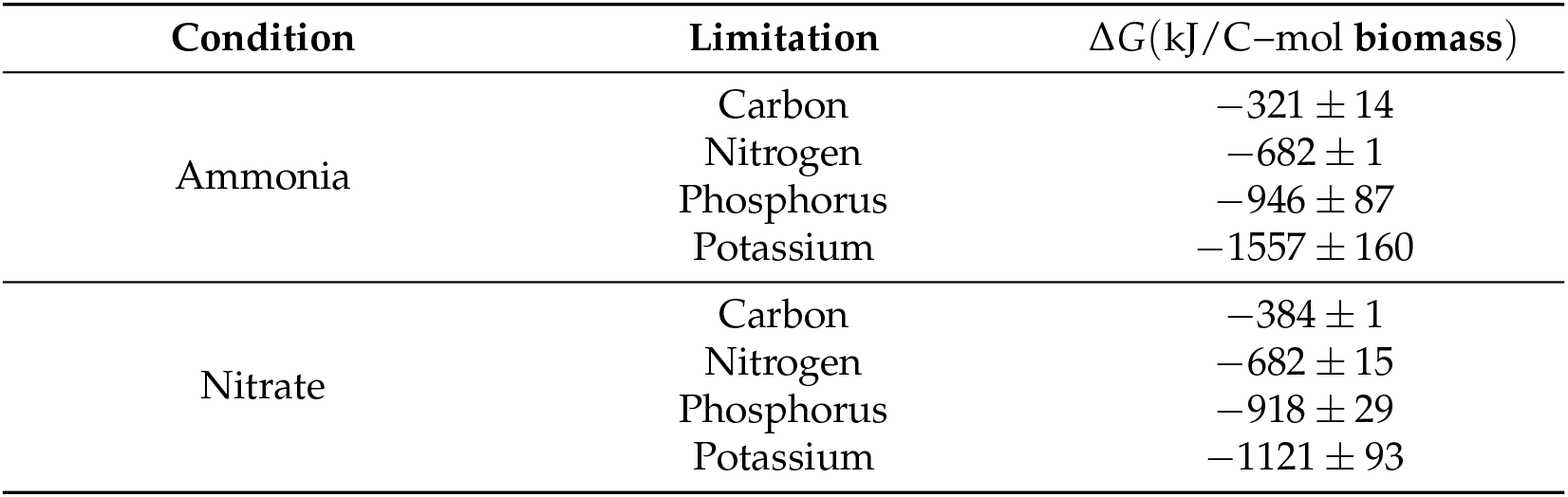
Summary of Fermentation Thermodynamics Under Different Continuous Growth Conditions and Nutrient Limitations of *Klebsiella pneumoniae* [12,35]. The table provides a schematic overview of the calculated overall thermodynamics of the growth reaction as described above across various nutrient-limited continuous growth conditions. The errors were estimated by correcting the discrepancy between fitted and measured fluxes either with biomass production fluxes or with respiration fluxes only.

| Condition | Limitation | $\Delta G$ (kJ/C–mol biomass) |
| --- | --- | --- |
| Ammonia | Carbon | $-321 \pm 14$ |
| | Nitrogen | $-682 \pm 1$ |
| | Phosphorus | $-946 \pm 87$ |
| | Potassium | $-1557 \pm 160$ |
| Nitrate | Carbon | $-384 \pm 1$ |
| | Nitrogen | $-682 \pm 15$ |
| | Phosphorus | $-918 \pm 29$ |
| | Potassium | $-1121 \pm 93$ |

**Figure 2.**
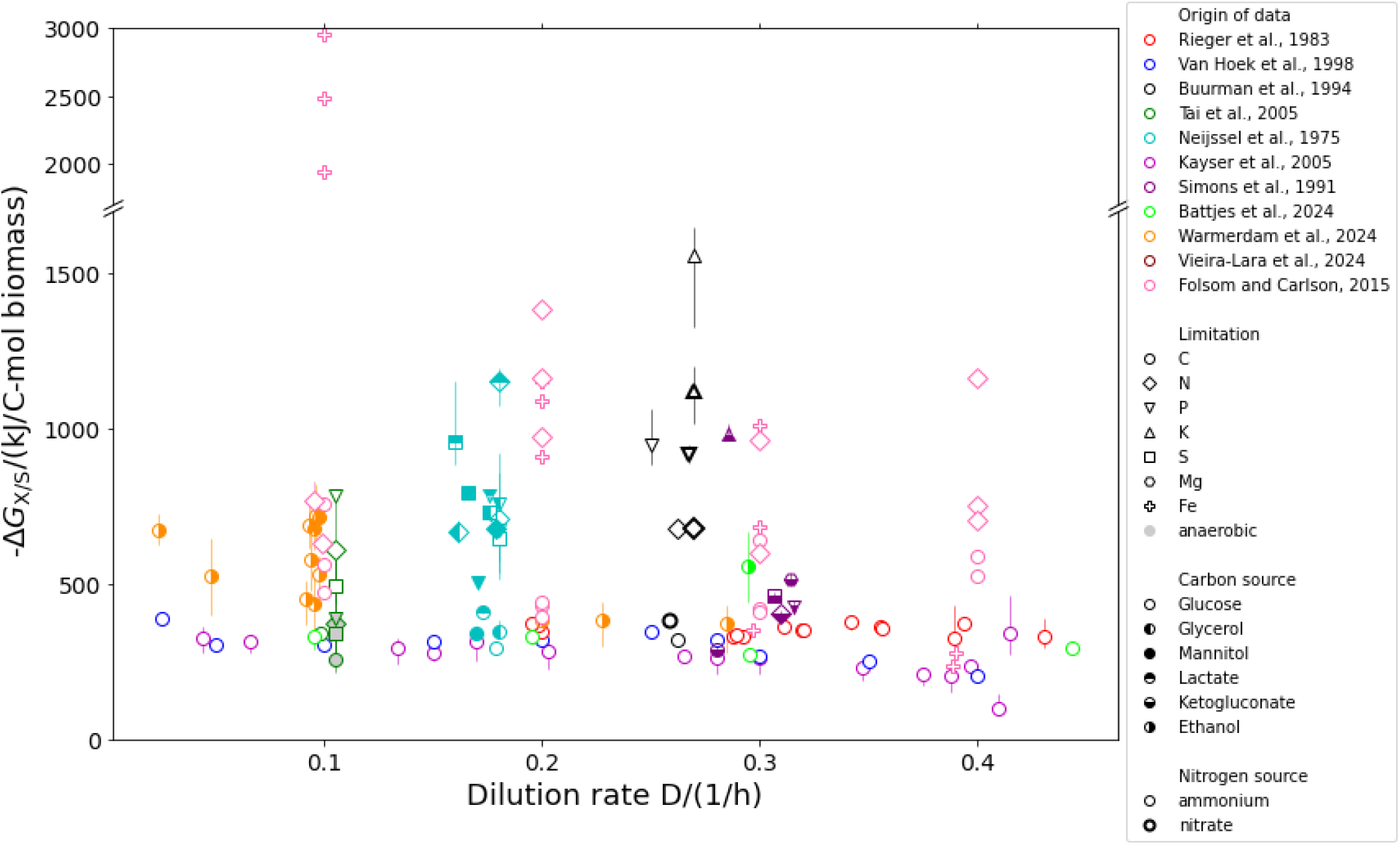
The calculated Gibbs free energy (Δ*G*_X/S_) of the Net Catabolic Reaction (NCR) during various growth conditions. All carbon-limited conditions seem to show limited growth rate dependent variation. It should be noted that the conditions above a dilution rate of *D* = 0.40 h^*−*1^ represent conditions that are close to mimicking batch conditions in which the carbon concentrations are no longer limited and fermentation products are produced in significant quantities. The error bars are estimated as described in Methods by correcting mismatching carbon recovery rates by either using the ideal anabolic reaction or pure respiration. Data from Folson and Carlson [34] did not include gas exchange rates, therefore the carbon recovery coefficient was not known and this approach this was not possible.

For the data on iron limitation [34], gas exchange was not measured but instead inferred. Therefore insufficient information is available for the above describe error estimation procedure. We decided to omit error bars for these data.

## 4. Results

In order to expand our current thermodynamic understanding of microbial growth to non-carbon limited conditions, we searched the literature for findings that address both carbon and non-carbon limited growth under comparable conditions. We limited our search to articles where carbon limited or anabolic limited continuous chemostat conditions were used and gas exchange, nutrient consumption and production formation rates were systematically measured to allow reconstruction of a balanced growth equation. In total, we selected 10 articles that fulfilled these criteria studying various yeasts including *Saccharomyces cerevisiae, Cyberlindnera jadinii* and *Pichia kluyveri* [18,19,25,35–38], *E. coli* [39], *Klebsiella aerogenes* [40] and *Klebsiella pneumoniae* [11,35]. In addition, we included one study on *E. coli*, in which gas exchange was not measured, to also include iron limitation in our analysis [34,41]. To obtain the thermodynamic characteristics, for each single experiment, we first determined a balanced macrochemical growth equation that best fits the experimentally determined exchange fluxes of nutrients and metabolic products and the set dilution (growth) rate. This fitting was necessary because of carbon recovery coefficients diverging from 100%. This equation was normalized to the production of 1 C-mol biomass, and the thermodynamic characteristics are calculated as described above (see Methods and Box 1). Our strategy extends previous approaches [10,30] by providing an estimate of the thermodynamics of catabolism normalized per unit of biomass formed and including non-carbon or non-energy limitations in our research [8,30].

We start our analysis with the study of Tai et al. [25], because it contains a large data set of *S. cerevisiae* under both aerobic and anaerobic conditions and four nutrient limitations (carbon, nitrogen, phosphorus and sulfur). We observed that the Δ*G*_X/S_ is more negative under all non-C-limited conditions (i.e. nitrogen, phosphorus and sulfur limitations) than during C-limited growth (see Table 1) under both aerobic and anaerobic conditions. The data demonstrate that under nutrient stress (i.e. nitrogen, phosphorus or sulfur limitation), the yeast cells require significantly more catabolic energy to sustain biomass production both under aerobic and anaerobic conditions. Our results indicate a clear pattern of increased catabolic energy demand with nutrient limitations.

Secondly we analyzed the available *K. pneumoniae* data (see Table 2 and [12]), which included chemostat experiments for two different nitrogen sources (ammonia and nitrate) under glucose-, nitrogen-, phosphorus-, and potassium-limited conditions. Again, determining the catabolic Gibbs free energy Δ*G*_X/S_, we observe that under nitrogen, phosphorus, and potassium limitation these values were consistently more negative than under carbon limitation.

To confirm these initial observations, we also included the other relevant articles that studied microbial growth under various nutrient limitations. Again, we applied the same strategy as above and determined the catabolic Gibbs free energy Δ*G*_X/S_ for every single reported chemostat experiment. This allowed us to use a broader dataset encompassing multiple microbial species, a variety of nutrient limitations (carbon, sulfate, nitrogen, phosphate, potassium, iron), and diverse carbon sources (glucose, ethanol, glycerol, mannitol, lactate, ketoglutarate). The entirety of the analyzed data is summarized in Figure 2.

Interestingly, first focusing on carbon limited conditions only (circles in Figure 2), reveals that despite variations in microbial species, media composition (all chemically defined), and dilution rates, the catabolic Gibbs free energy Δ*G*_X/S_ appears rather independent of organism, growth rate and type of catabolic carbon source used. It is noteworthy that both Crabtree positive (light green circles) and Crabtree negative (dark green circles) yeast strains showed a similar Δ*G*_X/S_ at all dilution rates, despite the drastically different catabolic routes and the concomitantly different Gibbs free energy yield per carbon mole substrate. Further partitioning Δ*G*_X/S_ into contributions from respiratory and nonrespiratory catabolism revealed that the respiratory component declines sharply with increasing growth rate, yet the total Δ*G*_X/S_ remains rather constant (see Figure 3). Notably, carbon limited growth on ethanol seemed to result in a slightly more negative Δ*G*_X/S_ of the NCR (see orange circles in Figure 2 and [36]).

**Figure 3.**
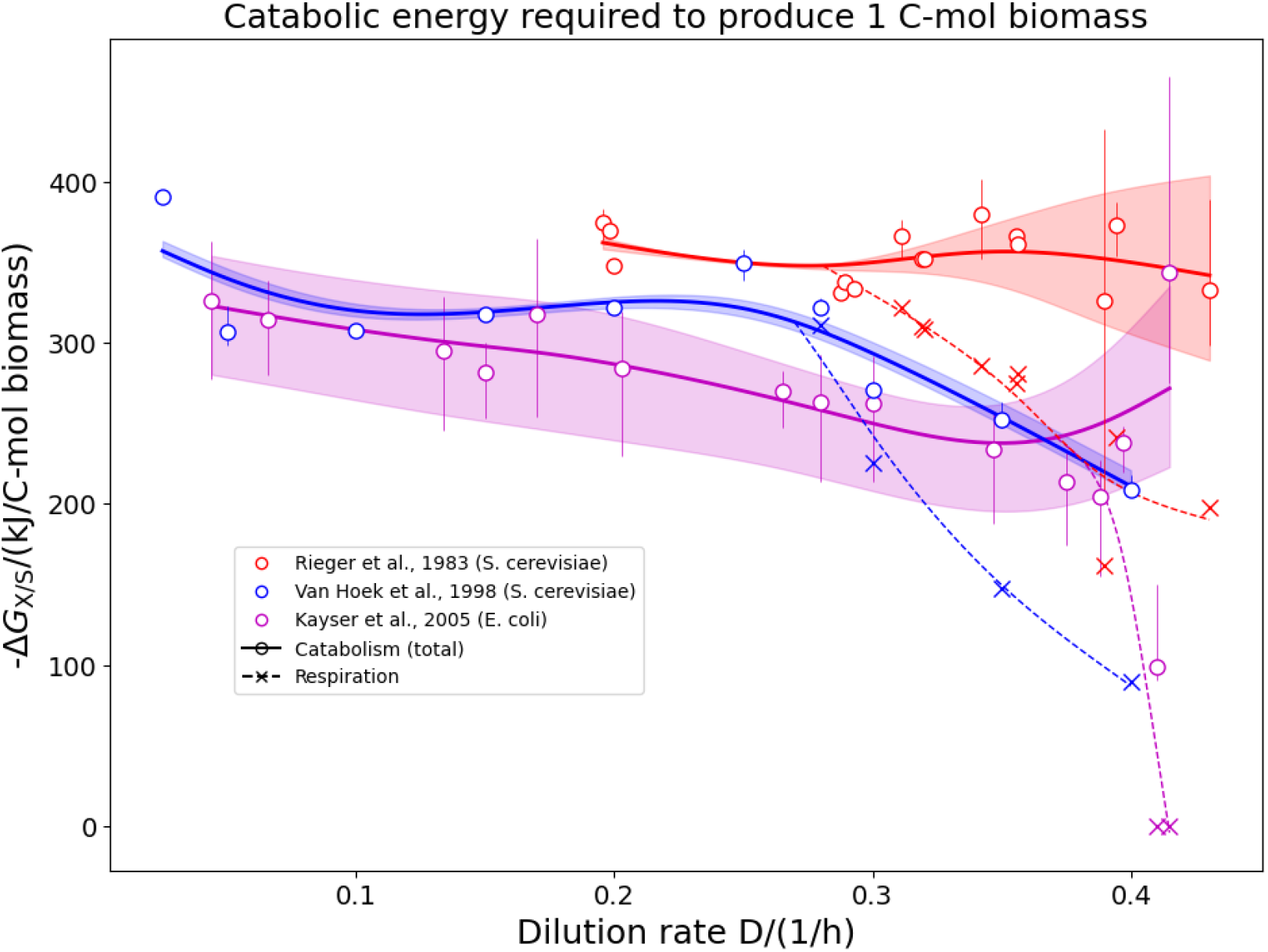
Calculated Gibbs free energy change (Δ*G*_X/S_) of the Net Catabolic Reaction (NCR) under carbon-limited conditions, showing both the total free energy for catabolism and the contribution from the respiratory component.

In contrast, under anabolic nutrient limitations (e.g., sulfur, nitrogen, phosphorus), the Δ*G*_X/S_ of the NCR was consistently more negative than for carbon limited conditions. This pattern held true across all examined organisms and conditions, indicating that anabolic limitations may universally impose a greater thermodynamic demand on catabolism than carbon limitation.

## 5. Discussion

Our analysis of nutrient-limited microbial growth provides additional insights on top of existing literature that studied the trade-offs between proteomic investment and thermodynamic efficiency under various growth rates under continuous C-limited growth conditions. There, it was consistently observed that under high growth rates catabolic fermentation products are excreted. This entails that the energetic substrate use efficiency decreases, because less Δ*G* is obtained per C-mol substrate, while simultaneously the yield decreases, and more substrate is consumed per C-mol biomass produced. This behavior is commonly explained [6,9,14] by arguing that the less efficient catabolic routes require fewer proteins but display a higher rate (*k*_cat_) and that, therefore, the same ATP production rate can be achieved with a smaller proteome fraction, leaving more space for anabolic proteins, in particular the ribosome. We think this hypothesis still holds, yet we cannot apply it to non-carbon limited conditions given that (1) under all non-C-limited growth conditions the Δ*G*_X/S_ of the overall growth equation is significantly more negative than under C-limited growth conditions for all microorganisms studied at similar growth rates; (2) the Δ*G*_X/S_ of the NCR is similar under both aerobic and anaerobic conditions (see dark green circles in Figure 2 and Table 1); (3) the effect is observed independent of the growth rate or carbon limitation used in the experimental setup. Moreover, it should be noted that knowledge of the catabolic equation alone does not allow discriminating between different catabolic pathways, and therefore also does not provide quantitative insight into the actual qATP. For example, both the ED and the EMP pathways may result in metabolizing one mole glucose into two moles pyruvate, while in the former case one mole of ATP was produced and in the latter two. Using the approach described here omits the use of any assumptions of catabolic pathways used and ATP produced and thereby circumvents this challenge. In fact, if we consider two organisms, one of which uses the ED and the other the EMP pathway, but which are otherwise identical and grow at identical rates, then the ED-using organism would need a catabolic flux twice as fast as the EMP-using organism to provide the same amount of ATP per time. With our approach, we would observe that the ED-using organism requires more free energy from catabolism to produce 1 C-mol of biomass. However, whether this apparent increased energy demand is actually caused by the usage of a lower yield pathway, or whether other energetic demands (such as caused by nutrient scarcity or external stresses) are present, cannot be discriminated from knowledge of the net catabolic reaction alone.

A remarkable observation of our analysis is that for C-limited conditions the catabolic Gibbs free energy, Δ*G*_X/S_, per C-mol biomass produced remains rather constant with values around Δ*G*_X/S_ = 300 … 400 kJ mol^*−*1^ (all circles in Figure 2). This value is relatively constant, irrespective of the organism investigated (*S. cerevisiae* in Rieger et al. [18] and van Hoek et al. [19], *E. coli* in Kayser et al. [39] and Folsom and Carlson [34], *K. pneumoniae* in Buurman et al. [12] and Simons et al. [35]), the carbon source used or the dilution rate applied. Most interestingly, this value is independent on whether ethanol (or any other overflow product) is released or not (compare the light and dark green circles in Figure 2 for data on the Crabtree positive yeast *P. kluyveri* and the Crabtree negative yeast *S. cerevisiae*, respectively, and see Figure 3 for three classical overflow experiments). This clearly indicates that under C-limited conditions, despite a rather drastic shift in metabolic pathways, the overall energetic requirement to produce 1 C-mol of biomass is rather independent on the growth rate. A slight increase of Δ*G*_X/S_ is visible for the yeast strains grown on ethanol (yellow points in Fig. 2, [36]) at low growth rates (≲ 0.1 h^*−*1^). However, for faster growth rates the Δ*G*_X/S_-value is no longer increased. Whereas proteome allocation principles may indeed explain the shift from respiratory to fermentative pathways, it does not make a statement regarding the overall catabolic energy requirement for growth.

To understand the above phenomena that the net catabolic Gibbs free energy Δ*G*_X/S_ is systematically more negative for limitations other than carbon, we think it relevant to highlight that during nutrient limitation, importing the limiting nutrient is expected to result in a more positive Δ*G* compared to conditions where nutrients are abundant as the concentration gradient from extracellular to intracellular space is higher during nutrient limitations. This more positive Δ*G* of nutrient uptake will add to the overall positive Δ*G* of biomass formation where disordered nutrients are concentrated and combined into a structured living cell [28]. This effect is not included in our calculations given that no information is available on the extracellular concentrations of the limiting nutrients.

It is unclear what the cause is of the more negative Δ*G*_X/S_ of the NCR under the various anabolic nutrient-limited growth conditions. This can either be caused by energy dissipation reactions or by overall changes in the catabolic pathways used, and possibly both. If one understands a growing microbe as an energy converter (see e.g. [29]), in which the negative Δ*G* of catabolism drives the (slightly positive) Δ*G* of anabolism, increasing the negative catabolic Δ*G* in non-carbon limited conditions makes intuitive sense. Nutrient limitations increase the energetic requirements to obtain these nutrients, e.g. by using active transporters or expressing extracellular pathways to capture sparse nutrients. Hence, an increased driving force of catabolism seems necessary to overcome the increased energy barrier of the driven anabolism. In the following, we postulate three, non-mutual exclusive, hypotheses on the mechanisms underlying a competitive advantage for the more negative Δ*G* under anabolic nutrient limitations:

1. **Proteome Allocation Hypothesis:** The more negative Δ*G* of the NCR under nutrient-limited conditions is a result of the preferential use of catabolic pathways with a lower ATP yield per substrate and thus a more negative Δ*G* (normalized to carbon mole new biomass formed). These pathways require fewer proteomic resources for ATP generation [3,14,16], freeing up a larger fraction of the proteome for the expression of nutrient transport systems and ribosomes. This redistribution could confer a selective advantage by enhancing the uptake of the limiting nutrient.
2. **Coupled Transport Contribution Hypothesis:** The more negative Δ*G* of the NCR may in part stem from the increased reliance on ATP-coupled or energetically driven transport mechanisms for nutrient uptake under limitation. When nutrients are scarce, cells may increasingly use active transporters that directly couple ATP hydrolysis to substrate import. While this incurs an energetic cost, it renders the overall nutrient uptake process thermodynamically more favorable and may contribute to the net negative Δ*G* of catabolism required to support growth. This strategy could enhance substrate import efficiency under nutrient limitation, thereby offering a selective advantage despite the higher energetic investment.
3. **Bioenergetic Efficiency Hypothesis:** The use of catabolic pathways with a more negative Δ*G* may also lead to increased cellular energy states, such as higher ATP/ADP ratios or higher membrane potentials. We hypothesize that this bioenergetic enhancement could improve the functionality of transporters that rely on ATP hydrolysis or membrane potential, thus supporting faster nutrient uptake under the anabolic limiting conditions discussed in this perspective. This could help overcome the additional thermodynamic burden of transporting the low concentration extracellular limiting nutrient to the highly concentrated intracellular environment of this nutrient.

All three strategies may contribute to the observed changes in overall Δ*G* of the NCR and it will prove challenging scientifically to disprove one or the other. In fact, all of the proposed mechanisms will result in an more negative catabolic Gibbs free energy Δ*G*_X/S_, and may therefore contribute to energetically and kinetically overcome nutrient limitations. Which of these mechanisms are actually active can be studied by detailed pathway analysis using stable isotope ^13^C labeling experiments that allow discrimination of the specific pathways used during specific conditions [42,43]. This may still however be challenging given that some of the microorganisms analyzed in this study show a branched respiratory chain that includes respiratory enzymes that differ in the number of protons translocated per oxygen reduced. Examples are the NDHII complex and cytochrome bd-I which both show a lower proton translocation than NDHI and Cytochrome bo, yet the *k*_cat_-values of these enzyme are significantly higher than their high efficiency counterparts [15,44–46]. It would therefore be relevant to further explore the catabolic routes and proteome resource allocation dynamics under anabolic nutrient limitations to provide further insights into the potential cause of the changes in Δ*G* of the NCR. Indications can be found in literature that this indeed may play a role as for *E. coli* the cytochrome bd-II complex shows increased expression under phosphate-limited conditions [26,47], implying that under phosphate limited continuous growth this catabolic pathway may play a role in the changes in overall Δ*G*_X/S_.

In general it is observed that although there is a strong shift in catabolic pathways used [6,18,21,39] the overall Δ*G* of the NCR does not show a significant change (see Figure 2) when switching from C-limited growth conditions to non C-limited growth conditions (i.e. batch growth). Although the Δ*G* per calculated ATP produced may change when such pathways are used (see Table 3 for some standard example calculations), it should be noted that such calculations assume specific catabolic pathways that may practically vary given the flexibility microorganisms tend to have in the respiratory chain (and therefore the P/O ratio of respiration) and in the used phosphorylation pathways. Therefore ATP based calculations are very challenging and we therefore suggest use of the NCR to ensure correct interpretation of the thermodynamics of catabolism.

**Table 3.** The driving force for ATP formation during heterotrophic growth for different catabolic pathways. The calculation of the Δ*G* was performed using eQuilibrator [9]. Values are given for biochemical standard conditions assuming a concentration of 1 mmol L^*−*1^.

| Organism | Reaction | $\Delta_r G^{1\text{mM}} (\text{kJ/C-mol ATP})$ | Growth |
| --- | --- | --- | --- |
| *L. lactis* | Glucose + 3 ADP + 3 $\text{P}_i \longrightarrow$ Acetate + Ethanol + 2 Formate + 3 ATP + 2 $\text{H}_2\text{O}$ | -43.8 | slow |
| *L. lactis* | Glucose + 2 ADP + 2 $\text{P}_i \longrightarrow$ 2 Lactate + 2 ATP + 2 $\text{H}_2\text{O}$ | -59.4 | fast |
| *S. cerevisiae* | Glucose + 36 ADP + 36 $\text{P}_i$ + 6 $\text{O}_2 \longrightarrow$ 6 $\text{CO}_2$ + 36 ATP + 36 $\text{H}_2\text{O}$ | -38.2 | slow |
| *S. cerevisiae* | Glucose + 2 ADP + 2 $\text{P}_i \longrightarrow$ 2 Ethanol + 2 $\text{CO}_2$ + 2 ATP + 2 $\text{H}_2\text{O}$ | -93.9 | fast |
| *E. coli* | Glucose + 28 ADP + 28 $\text{P}_i$ + 6 $\text{O}_2 \longrightarrow$ 6 $\text{CO}_2$ + 28 ATP + 28 $\text{H}_2\text{O}$ | -61.6 | slow |
| *E. coli* | Glucose + 5 ADP + 5 $\text{P}_i$ + 2 $\text{O}_2 \longrightarrow$ 2 Acetate + 2 $\text{CO}_2$ + 5 ATP + 5 $\text{H}_2\text{O}$ | -198.7 | fast |

## 6. Conclusion

In this study, we have shown that a thermodynamics analysis of the Net Catabolic Reaction (NCR) provides a powerful framework for understanding the energetic strategies of microbial metabolism. The assumptions made to derive the NCR are universal and thus applicable to any carbon source and organism. By examining the Gibbs free energy changes associated with the overall NCR across various nutrient limitations and microorganisms, we reveal that microbes consistently use more free energy from catabolic pathways to produce biomass under anabolic nutrient limitations compared to carbon limited conditions. This thermodynamic shift is not merely a passive consequence of nutrient limitation but we hypothesize that this is an active cellular strategy to optimize proteome resource allocation and uptake rate, or the apparent *K*_M_ of the cell for the limiting nutrient.

Our findings extend existing theories on the thermodynamic interpretation of carbon-limited growth at various growth rates to other nutrient limitations, such as nitrogen, sulfate, potassium and phosphate. The observations suggest that thermodynamic optimization of the catabolic pathways used under such anabolic nutrient limitations is a general principle shaping microbial physiology. These insights pave the way for a broader understanding of how thermodynamic and proteomic constraints co-determine microbial fitness in specific nutrient-limited environments. Future work combining experimental flux measurements, proteomic profiling, and thermodynamic modelling under defined nutrient limitations will be crucial to validate and further refine these hypotheses. Ultimately, integrating thermodynamics into systems biology not only deepens our comprehension of microbial adaptation but also holds promise for the rational design of microbial strains for biotechnology and bioprocessing applications.

Understanding microbial thermodynamics has profound implications for bioproduction and industrial microbiology. It shows that some processes will inherently dissipate heat under certain conditions. A good example is the production of omega-3 which is done using various microorganisms under nitrogen limited conditions [48]. It would be relevant to study whether heat dissipation can be minimized by a better understanding of the causes of the more negative Gibbs free energy, Δ*G*_X/S_, of the NCR. Moreover, expanding the current framework beyond hydrocarbon-based catabolism to include alternative electron donors, such as ammonia, would enhance its applicability. Extending the theory to ammonia oxidation is conceptually straightforward, as the separation of the macrochemical growth equation into anabolic and catabolic components remains valid. However, this would require experimental determination of exchange rates for nitrogen-containing metabolites. Future research should explore the integration of thermodynamic modeling with synthetic biology approaches, particularly in the context of precision fermentation and metabolic engineering.

## Glossary

**Symbol Meaning**

Δ*G*: Gibbs free energy change of a reaction (kJ mol^−1^ or kJ (C−mol biomass)^−1^); indicates thermodynamic favorability
NCR: Net Catabolic Reaction; calculated Δ*G* of catabolism normalized per c-mole of biomass formed
*Y*_ATP_: Yield of ATP per mole of substrate consumed
qATP: Specific ATP production rate (mol ATP per mol biomass per hour)
*k*_cat_: Turnover number of an enzyme; number of substrate molecules converted per enzyme per second
*K*_M_: Michaelis constant; substrate concentration at which reaction rate is half of its maximum
*γ*_*X*_: Degree of reduction of biomass
*γ*_*S*_: Degree of reduction of substrate
Δ_*r*_*G*^*1mM*^*/ATP*: Gibbs free energy change per mole of ATP formed under biochemical standard conditions (1 mM concentrations)
C-mol: Carbon mole; a unit representing the amount of carbon in biomass or substrate
P/O ratio: Phosphate/Oxygen ratio; number of ATP molecules synthesized per atom of oxygen reduced during oxidative phosphorylation; this reflects respiratory chain efficiency
EMP: Embden-Meyerhof-Parnas pathway; classical glycolysis pathway
ED: Entner-Doudoroff pathway; alternative glycolysis pathway with lower protein cost

## Author Contributions

Conceptualization, M.B. and O.E.; methodology, M.B. and O.E.; software, O.E.; validation, M.B. and O.E.; writing—original draft preparation, M.B.; writing—review and editing, M.B. and O.E.; All authors have read and agreed to the published version of the manuscript.

## Institutional Review Board Statement

Not applicable

## Informed Consent Statement

Not applicable

## Data Availability Statement

All code developed for this project is found at https://gitlab.com/qtb-hhu/thermodynamics-task-force/2025-catabolic-deltag-for-various-limitations. Files and instructions are given how to reproduce the results and figures in this publication.

## Disclaimer/Publisher’s Note

The statements, opinions and data contained in all publications are solely those of the individual author(s) and contributor(s) and not of MDPI and/or the editor(s). MDPI and/or the editor(s) disclaim responsibility for any injury to people or property resulting from any ideas, methods, instructions or products referred to in the content.

